# Transgenerational effects of perinatal cannabis exposure on female reproductive parameters in mice

**DOI:** 10.1101/2025.02.24.639897

**Authors:** Mingxin Shi, Yeongseok Oh, Debra A. Mitchell, James A. MacLean, Ryan J. McLaughlin, Kanako Hayashi

## Abstract

The use of cannabis during pregnancy and nursing is a growing public health concern, and the multigenerational impacts of perinatal cannabis exposure remain largely unknown. To address this knowledge gap, we sought to examine the long-term consequences of perinatal cannabis use on reproductive function and how it might impact subsequent generations. Pregnant female mice were exposed to control vehicle or cannabis extract [25, 100, or 200 mg/ml Δ9-tetrahydrocannabinol (THC) in the cannabis extract] from gestational day 1 to postnatal day 21 (twice/day), encompassing the duration of pregnancy through weaning. Based on plasma THC concentrations in F0 females, we chose 100 and 200 mg/ml THC in the cannabis extract for subsequent studies. The selected doses and exposure conditions did not disrupt pregnancy or nursing in F0 females. Pregnancy and neonatal outcomes, including gestational length, litter size, and sexual ratio, were not affected by cannabis exposure. However, cannabis-exposed neonatal F1 pups were smaller. Cannabis exposure delayed vaginal opening as a sign of puberty onset and disrupted estrous cyclicity in F1 females. However, its effects were minor in F2 and F3 females. F1-F3 females showed no abnormal ovarian and uterine histology or plasma estradiol-17β levels and could produce normal offspring without pregnancy issues. These results suggest that the hypothalamus and pituitary are likely perturbed by perinatal cannabis exposure, and the early hypothalamus-pituitary-ovarian axis is disrupted in F1 females. However, they are not sufficient to compromise adult reproductive function. The present results indicate limited transgenerational effects of perinatal cannabis exposure on female reproductive parameters.

## INTRODUCTION

Cannabis is considered the most widely used recreational drug in the United States (US) and all over the world. The National Survey on Drug Use and Health (NSDUH) reported that over 48 million Americans aged 12 and older used cannabis in the past year ((SAMHSA) 2020). This number has been increasing due to the expanding legalization of cannabis consumption in the US. Since 2012, a total of 24 states, Washington D.C., and three territories have legalized the recreational sale and use of cannabis. The prevalence of cannabis use has increased explicitly among reproductive-aged people and was heightened during the COVID-19 pandemic (Imtiaz et al. 2021; Young-Wolff et al. 2021). Cannabis use in pregnancy also increased by 7% in 2017 (Volkow et al. 2019) and up to 25% during the COVID-19 pandemic (Young-Wolff et al. 2021). Simultaneously, following the increase in legalized recreational cannabis use, higher potency strains of cannabis plants have emerged over the past two decades (ElSohly et al. 2016). Indeed, the average concentration of Δ^9^-tetrahydrocannabinol (THC), the primary psychoactive component of cannabis, has increased from 9% to 17% in the past 10 years. The ratio of THC to cannabidiol (CBD), the primary non-psychoactive compound, has also risen substantially from 23 (THC/CBD) in 2008 to 104 in 2017 (Chandra et al. 2019). Unsurprisingly, adverse health outcomes associated with cannabis consumption have also been increasing (Belladelli et al. 2021). However, the NSDUH has reported that approximately 70% of pregnant and non-pregnant women believe there to be only a slight risk or no risk of harm from using cannabis once or twice a week during pregnancy (Ko et al. 2015). The American College of Obstetricians and Gynecologists recommends that people abstain from using cannabis during pregnancy and breastfeeding due to the association between cannabis use and adverse pregnancy outcomes (American College of and Gynecologists Committee on Obstetric 2015). As THC can readily cross the placenta (Blackard and Tennes 1984), in utero cannabis exposure has been reported to be associated with stillbirth, preterm birth, fetal growth restriction, and/or impaired fetal neural development (Conner et al. 2016; Gunn et al. 2016; Lo et al. 2024; Marchand et al. 2022).

The endocannabinoid system (ECS) is present in many cell types and tissues to regulate female reproductive systems (Di Blasio et al. 2013; Walker et al. 2019). Small natural lipids endocannabinoids, such as N-arachidonoylethanolamine (anandamide; AEA) and 2-arachidonoylglycerol (2-AG), are essential regulators for hypothalamic-pituitary-ovarian (HPO) axis, folliculogenesis, oocyte maturation, ovulation, fertilization, implantation, decidualization, and placentation (Walker et al. 2019) and their actions are mediated by type-1 and type-2 cannabinoid receptors (CB1 and CB2, respectively). Exogenous THC exerts its effects via partial agonism of the same receptors and, when bound, can interfere with endocannabinoid regulation of reproductive function (Pertwee et al. 2010). In fact, THC can suppress female fertility by disrupting hypothalamic gonadotropin-releasing hormone (GnRH) secretion (Brents 2016; Chakravarty et al. 1979) and preventing follicle-stimulating hormone (FSH), luteinizing hormone (LH), and prolactin release from the anterior pituitary (Brents 2016; Mendelson et al. 1985), leading to reduced estrogen and progesterone production and anovulatory menstrual cycles (Brents 2016). Bolus administration of THC causes disruption of estrous cycles in rats (Dhawan and Sharma 2003). THC interferes with the cellular function of oocytes and embryos (Dubovis and Muneyyirci-Delale 2020). Furthermore, strong correlations exist between cannabis use by pregnant women and adverse obstetrical, pregnancy, and neonatal outcomes (Dubovis and Muneyyirci-Delale 2020; Fonseca and Rebelo 2021). Parental cannabis use increases the risks of congenital malformation (Reece and Hulse 2019; van Gelder et al. 2014; van Gelder et al. 2010; Weinsheimer et al. 2008; Williams et al. 2004), psychosis (Day et al. 2015; Fine et al. 2019; Grant et al. 2018; Zammit et al. 2009), and delinquent behavior (Day et al. 2011; Goldschmidt et al. 2000; Porath and Fried 2005; Sonon et al. 2015) among in utero exposed offspring. Recent studies further highlight the ability of THC exposure to induce significant changes in DNA methylomes in spermatozoa (Murphy et al. 2018). However, the long-term consequences of cannabis use on female reproductive function and how it might impact the next generation have not been examined.

In the present study, we examined the multi- and trans-generational effects of cannabis vapor exposure on female reproductive function. Intrapulmonary delivery is the most common route of cannabis administration in humans (Baggio et al. 2014; Lee et al. 2016), whereas purified THC, CBD, and/or synthetic CB1 agonists have been dosed to animals via either intraperitoneal injection (i.p.), intravenous exposure (i.v.), or oral gavage (p.o.) to assess the physiological and/or toxicological effects of cannabis (Carvalho et al. 2020). In order to understand the multi- and trans-generational effects of cannabis exposure on female reproductive function, the present study was performed using a model of vaporized cannabis exposure, whereby adult pregnant and nursing females were exposed to cannabis extract to assess the toxicological effects of cannabis in F1, F2, and F3 females. We report that cannabis vapor exposure to pregnant and nursing females delayed vaginal opening and/or impaired estrous cyclicity in F1 and F2 offspring. However, perinatal cannabis exposure did not disrupt pregnancy and nursing in F0 females, adult reproductive function, or pregnancy and neonatal outcomes in F1, F2, and F3 females. Our results suggest that vaporized cannabis exposure directly affects the early stages of female reproductive parameters in F1 females. Conversely, our data indicate limited multi- and trans-generational effects of in utero and nursing cannabis exposure on female reproductive function.

## MATERIALS AND METHODS

### Animals

CD-1 mice were purchased from Inotiv and housed in an environment-controlled animal facility (12:12 light-dark cycle) with ad libitum access to food and water. All animal experiments were performed at Washington State University according to the NIH guidelines for the care and use of laboratory animals (protocol #7108).

### Drugs

Whole-plant cannabis extract was obtained from the National Institute on Drug Abuse (NIDA) Drug Supply Program. The raw extract contained 69.8% THC, 0.89% tetrahydrocannabivarin (THCV), 0.83% cannabichromene (CBC), 2.69% cannabigerol (CBG), 1.51% cannabinol (CBN), and 0.73% Δ8 tetrahydrocannabinol (Δ8 THC). CBD concentration is below the threshold of detection. The total terpene concentration present in the extract is 1.35%. The extract was prepared based on the final THC potency at 25, 100, or 200 mg/ml THC concentration by dissolving raw cannabis extract into vehicle control of polyethylene glycol (PEG) 400 under continuous stirring at 60 °C as described previously (Freels et al. 2020).

### Study Design

To investigate the multi- and trans-generational effects of in utero and nursing exposure to cannabis, outbred CD-1 mice were used because they are considered optimal for toxicological studies as they are outbred and, thus, a closer model to human populations. A study design schematic with a mating scheme is shown in Figure. 1A. Adult female mice (8-10 weeks old) were mated with males. The presence of a vaginal plug was defined as gestational day (GD) 1, and females were randomly assigned to control vehicle or cannabis groups (25, 100, or 200 mg/ml THC = 35.8, 143.3, or 286.5 ng/ml cannabis extract, respectively, n=10/group). F0 females were exposed to vaporized control vehicle or cannabis extract (25, 100, or 200 mg/ml THC), receiving 3 sec puffs of vapor every 2 min for 30 min twice daily beginning at 8 am and 5 pm from GD1 to postnatal day (PND) 21 using the custom-made vapor chamber (La Jolla Alcohol Research, Inc.). This window covered the entire pregnancy to weaning. Because dams were separated from pups during exposure, exposure to offspring could only be via the placenta or milk. To avoid complications around birth, F0 females were not exposed between GD19 at 5 pm, a day before parturition, and 4 days after birth. Mice delivered from F0 females were labeled as the F1. To generate F2, F1 females were bred to drug-naïve CD-1 males, which was repeated in the F3 generation. The remaining F1, F2, and F3 offspring from these litters were used for the studies.

**Figure 1.**
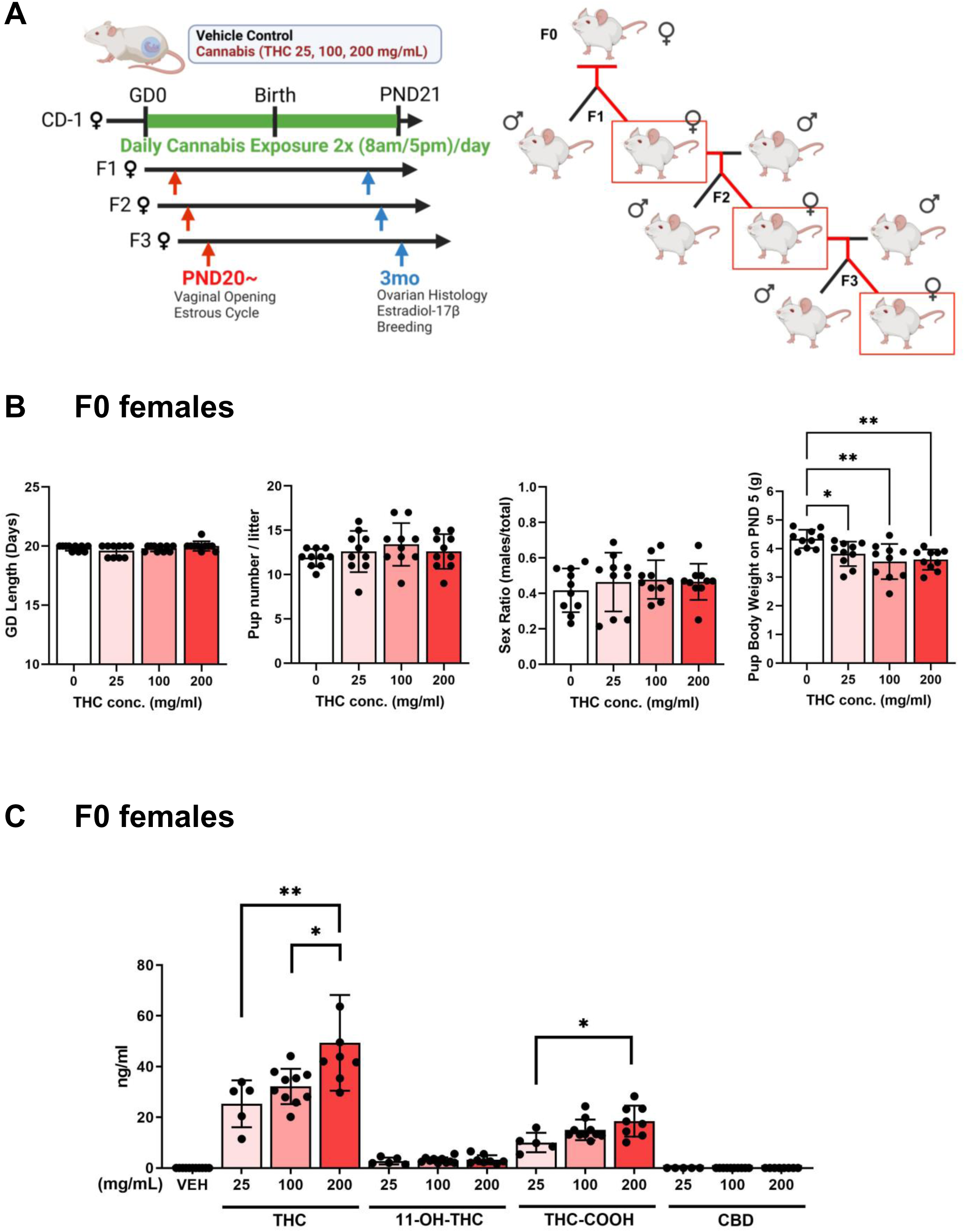
Schematic diagram of the experimental design. (A) Adult CD-1 female mice (F0) were mated with CD-1 males and were exposed to control vehicle or cannabis extract (25, 100, or 200 mg/ml THC, n=10/group) 2x/day from gestational day (GD) 1 to postnatal day (PND) 21. Mice delivered from F0 females were labeled as the F1. To generate F2, F1 females were bred to drug-naïve CD-1 males, which was repeated in the F3 generation. Created with BioRender.com (B) gestational length of F0 females, pup numbers at birth, sex ratio of pups, and pup body weights on PND5 are shown. (C) plasma THC, 11-OH-THC, THC-COOH, and CBD in F0 females were analyzed by mass spectrometry within 10 min after the last cannabis exposure (n=5-10/group). \**P* < 0.05, \*\**P* < 0.01.

Starting at PND20, female pups from each dam were examined for vaginal openings daily by visually examining the vulva, indicating the onset of puberty following our previous studies (Shi et al. 2017; Shi et al. 2019a; Shi et al. 2019b). Once vaginal opening occurred, estrous cyclicity was evaluated by examining vaginal smears daily for 40 days (Shi et al. 2017; Shi et al. 2019a; Shi et al. 2019b). Female pups (1-2 females from each dam) were mated with drug-naïve CD-1 males and examined for a plug as a sign of successful mating. After a vaginal plug was observed, the female was separated from the male, and gestational length, pup numbers, and the sex ratio of pups were recorded. The remaining female pups, which were not for breeding study, were euthanized to collect blood for hormone assays and mass spectrometry and both ovary and uterus for histological analysis. The results obtained from pups from the same litter were averaged and used as one representative data point for statistical analysis.

### Blood Collection and Mass Spectrometry

F0 females were euthanized after the weaning of pups within 10 min from the last session for blood collection to analyze plasma THC and THC metabolite concentrations. Cannabinoids in plasma (THC, 11-OH-THC, THC-COOH, and CBD) were analyzed by the Washington State University Tissue Imaging, Metabolomics, and Proteomics Laboratory, as previously described (Britch et al. 2017) using a Synapt G2-S (Waters) mass spectrometer. Samples with <1ng/ml were considered below the detection limit and listed as a value of 0.

### ELISA

Plasma levels of estradiol-17β were measured by ELISA (501890, Cayman Chemical) following the manufacturer’s instructions.

### Statistical Analysis

Statistical analyses were performed using GraphPad Prism (version 9.5). Data were tested for normal distribution using the Shapiro-Wilk normality test. If data were normally distributed, one-way ANOVA followed by Tukey multiple comparison tests or an unpaired two- tailed t-test were used to analyze the differences among the groups or between the two groups, respectively. If data were not normally distributed, the Mann-Whitney or Kruskal-Wallis test was performed. A *P* value less than 0.05 was considered to be statistically significant.

## RESULTS

### Effects of perinatal cannabis exposure in F0 females

As many studies associate maternal cannabis use with preterm birth and stillbirth (Dubovis and Muneyyirci-Delale 2020), we first examined gestational length in F0 females, pup numbers at birth, sex ratio of pups, and pup body weights on PND5 (Figure 1B). Vapor exposure to cannabis with any doses during the entire pregnancy did not induce preterm birth or stillbirth and did not affect gestational length, pup numbers, and sex ratio. However, the body weights of F1 pups on PND5 were reduced in all cannabis groups, but there were no differences among the groups.

Plasma levels of THC and its metabolites, 11-OH-THC, and THC-COOH, as well as CBD, were analyzed in F0 females immediately after the last session of cannabis vapor exposure (Figure 1C). As expected, THC, 11-OH-THC, THC-COOH, and CBD were not detected or below the detection limit in the control group (Figure 1C). Thus, we only showed one vehicle control (VEH, value = 0). Plasma THC concentrations within 10 min after chronic cannabis exposure reached 25.3, 32.2, and 49.3 ng/ml from doses of 25, 100, and 200 mg/ml THC of cannabis extract, respectively. A dose of 200 mg/ml showed significantly higher THC in plasma than other groups. Plasma 11-OH-THC was at 2.8, 3.1, and 3.3 ng/ml from doses of 25, 100, and 200 mg/ml, respectively, and did not show any differences between groups. Plasma THC-COOH, which is a significant metabolite that has a longer half-life and is generally used for the detection of cannabis use in humans, was detected at 10.1, 15.1, and 18.5 ng/ml from doses of 25, 100, and 200 mg/ml, respectively. THC-COOH level in the group of 200 mg/ml THC was higher than in the group of 25 mg/ml THC. We confirmed undetectable levels of CBD in plasma, as CBD concentration is below the detection threshold in the cannabis extract that was used for the study. As doses of 25 and 100 mg/ml THC did not show any differences in the plasma concentrations of THC, 11-OH-THC, or THC-COOH, we chose 100 and 200 mg/ml THC groups for further analysis.

### Effects of perinatal cannabis exposure on reproductive function in F1 females

To investigate the direct effects of in utero and nursing cannabis vapor exposure on female reproduction in F1 offspring, we examined the presence of a vaginal opening day as an indicator of the age of puberty onset and estrous cyclicity following the vaginal opening (Figure 2). In F1 females, the vaginal opening was dose-dependently delayed in cannabis-exposed groups. In particular, the 200 mg/ml THC group showed delayed vaginal opening for approximately 10 days (∼PND 37.1) than the control group (∼PND 27.1) (Figure 2A left). The vaginal opening in the 100 mg/ml THC group was at ∼PND 31.1. When we examined estrous cyclicity for 40 days following the vaginal opening, control mice exhibited standard 4–6 days of the estrous cycle and a total of 6.1 cycles per 40 days (the first estrous to the next estrous was counted as one cycle). However, F1 females in the 100 or 200 mg/ml THC groups showed irregular estrous cyclicity with fewer cycles, a total of 5.0 or 3.3 cycles, respectively (Figure 2A middle). In association with irregular and fewer estrous cycles, each cycle was longer in the 100 (6.5 days) and 200 (8.9 days) mg/ml THC groups than the control group (5.2 days) (Figure 2A right). In contrast, body and ovarian weights and plasma estradiol-17β levels after examining estrous cyclicity for 40 days were not affected by cannabis exposure (Figure 2BC). Furthermore, we did not observe abnormal ovarian and uterine histology among the groups (Figure 3A). When F1 females were mated with drug-naïve males, all F1 females received successful pregnancies, and their gestational length, pup numbers, sex ratio of pups, and pup body weights on PND5 were not affected by cannabis exposure (Figure 3B).

**Figure 2.**
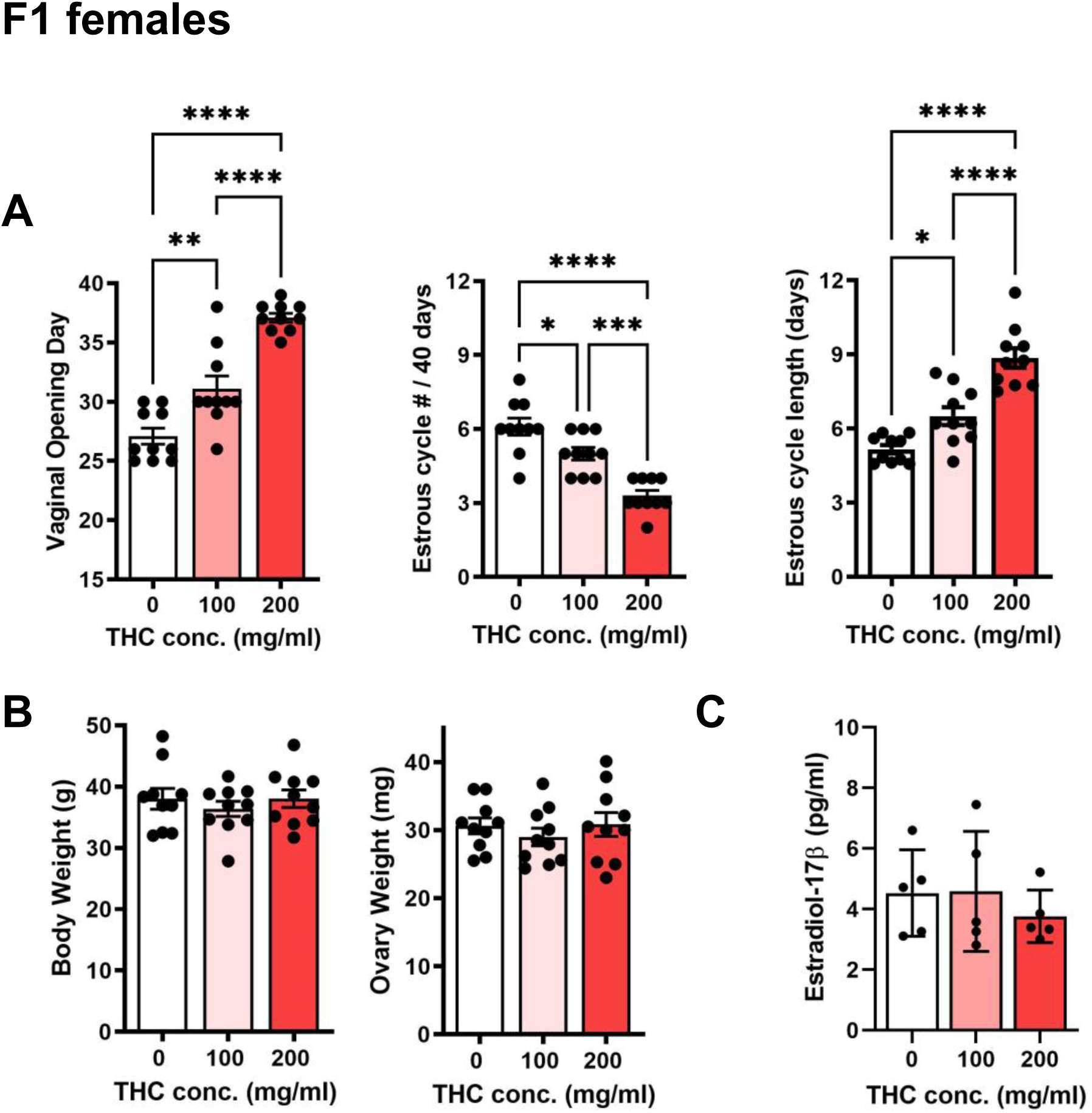
Effects of in utero and nursing cannabis vapor exposure on (A) vaginal opening day, estrous cycle number for 40 days after vaginal opening occurred, and estrous cycle length for 40 days (n=10/group) in F1 females. (B) Body and ovarian weight (n=10/group) and (C) plasma estradiol 17β (n=5/group) in F1 adult females at 3-mo of age. \**P* < 0.05, \*\**P* < 0.01, \*\*\**P* < 0.001, **** *P* < 0.0001.

**Figure 3.**
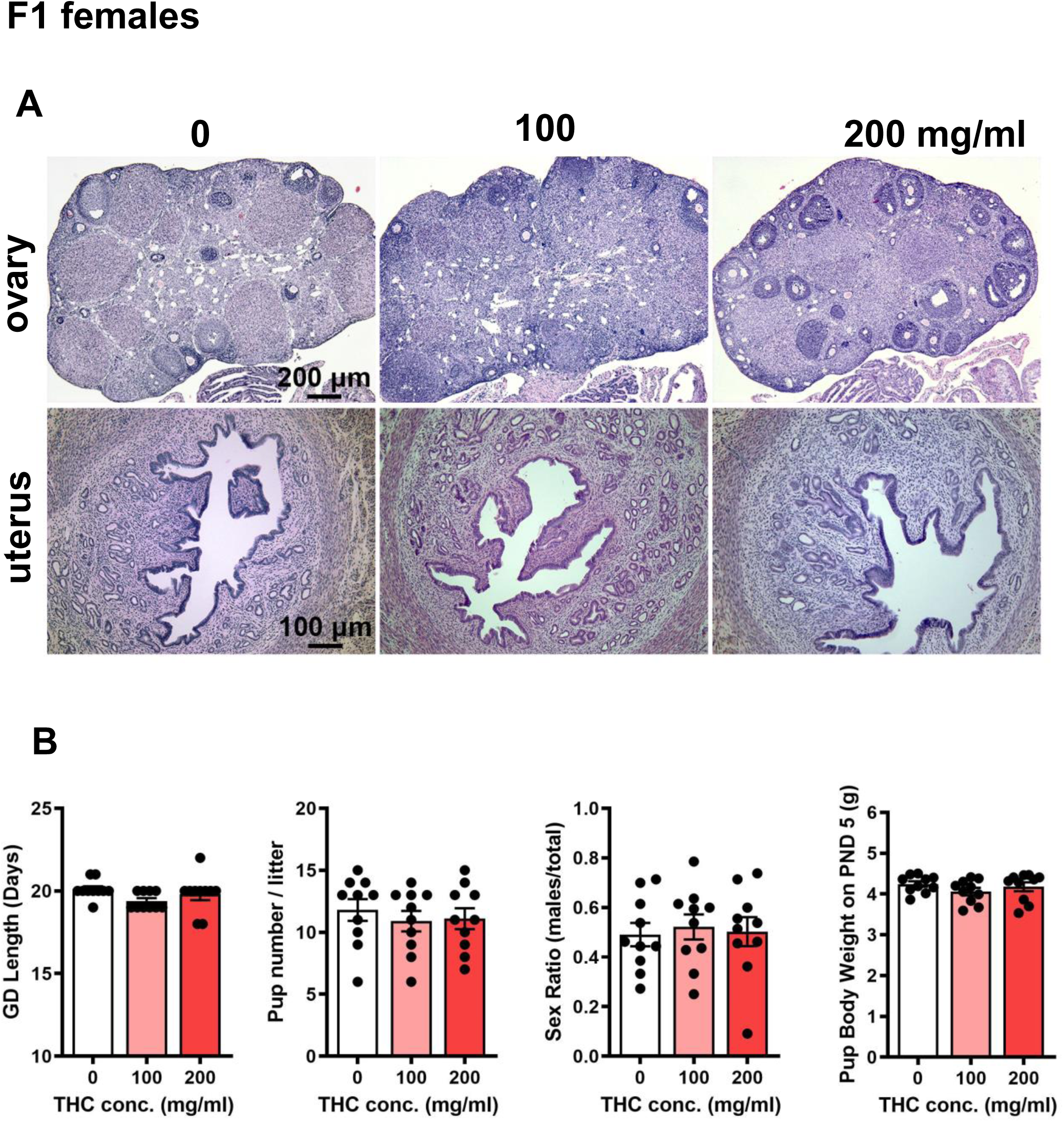
Effects of in utero and nursing cannabis vapor exposure on (A) ovarian and uterine histology and (B) gestational length of F1 females, pup numbers at birth, sex ratio of pups, and pup body weights on PND5 (n=10/group).

### Effects of perinatal cannabis exposure on reproductive function in F2 and F3 females

Although F1 females showed dose-dependent effects of delayed puberty and irregular estrous cyclicity, their adult reproductive parameters, such as successful pregnancy and pregnancy outcomes, were normal. Thus, we only examined the 200 mg/ml THC group in F2 and F3 females (Figures 4 and 5). In F2 females, 200 mg/ml THC did not alter or delay vaginal opening (Figure 4A left). At the same time, they still had irregular estrous cyclicity with fewer cycles at 5.6 cycles per 40 days, compared with 7.1 cycles in the control group (Figure 4A middle). The average estrous cycle length (6.0 days) in the 200 mg/ml THC group was longer than 4.9 days in the control group (Figure 4A right). However, body and ovarian weights and plasma estradiol-17β levels in F2 females were not affected by cannabis exposure (Figure 4 BC). We did not observe abnormal ovarian and uterine histology (Figure 4D). Furthermore, F2 females did not show any pregnancy issues, including gestational length, pup numbers, sex ratios of pups, and pup body weights on PND5 (Figure 4E).

**Figure 4.**
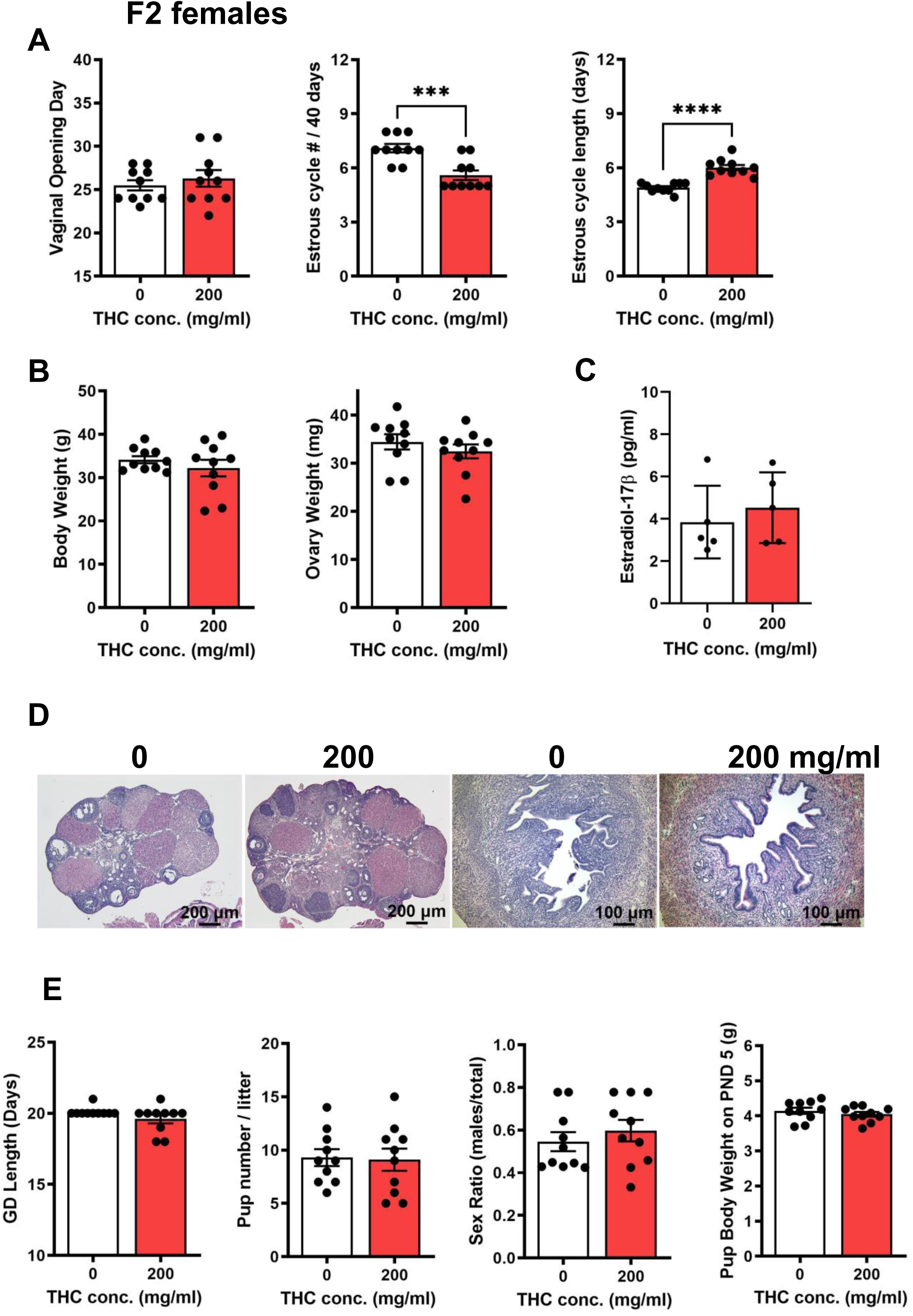
Effects of in utero and nursing cannabis vapor exposure on (A) vaginal opening day, estrous cycle number for 40 days after vaginal opening occurred, and estrous cycle length for 40 days (n=10/group) in F2 females.\*\*\**P* < 0.001, **** *P* < 0.0001. (B) Body and ovarian weight (n=10/group) and (C) plasma estradiol 17β (n=5/group) in F2 adult females at 3-mo of age. (D) ovarian and uterine histology, and (E) gestational length of F2 females, pup numbers at birth, sex ratio of pups, and pup body weights on PND5 (n=10/group).

**Figure 5.**
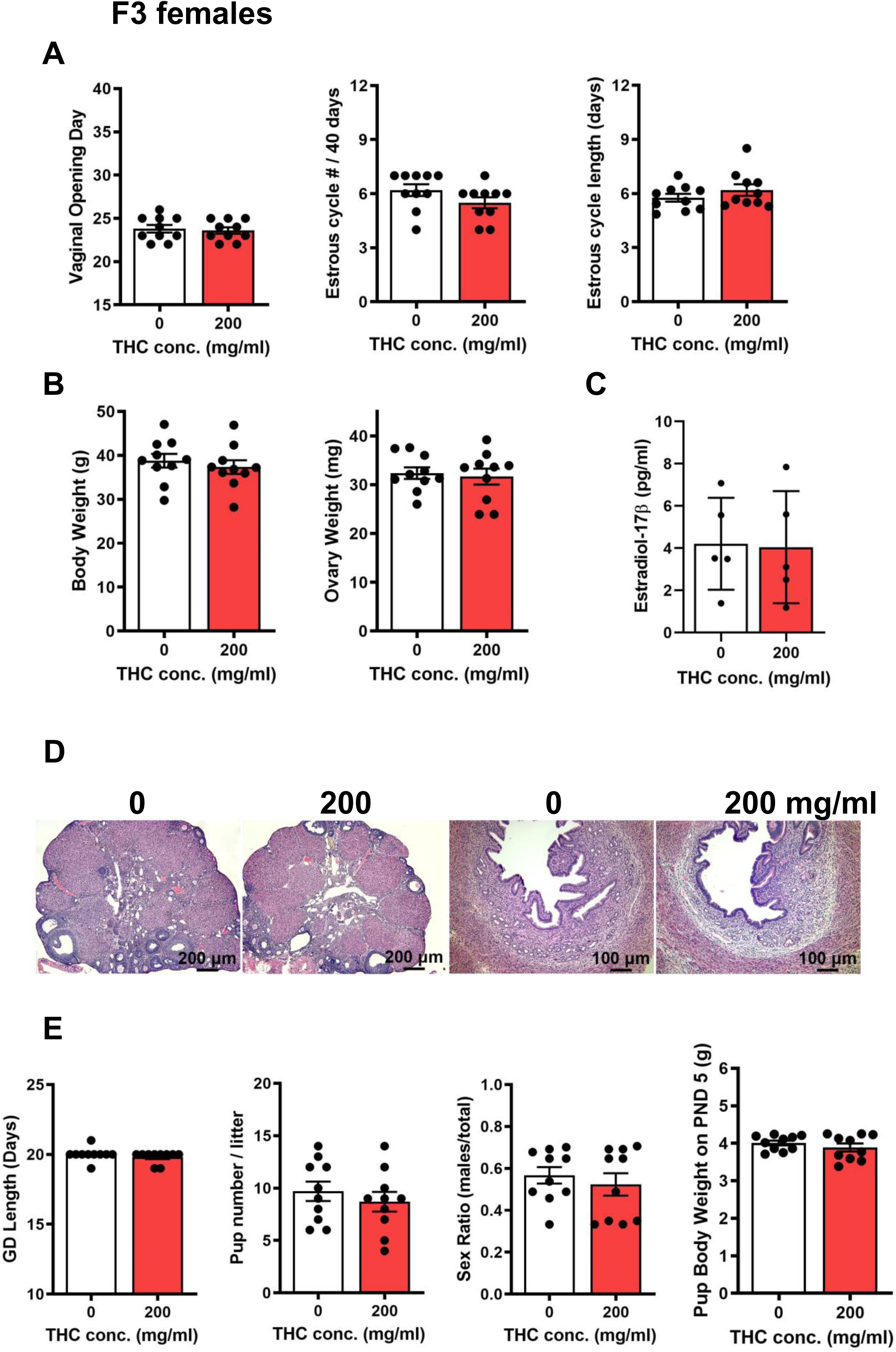
Effects of in utero and nursing cannabis vapor exposure on (A) vaginal opening day, estrous cycle number for 40 days after vaginal opening occurred, and estrous cycle length for 40 days (n=10/group) in F3 females. (B) Body and ovarian weight (n=10/group) and (C) plasma estradiol 17β (n=5/group) in F3 adult females at 3-mo of age. (D) ovarian and uterine histology, and (E) gestational length, pup numbers at birth, sex ratio of pups, and pup body weights on PND5 (n=10/group).

In F3 females, vaginal opening, estrous cyclicity, body and ovarian weights, and plasma estradiol-17β levels were no longer affected by F0 maternal cannabis exposure (Figure 5 ABC). We did not find any abnormal ovarian and uterine histology, gestational length, pup numbers, sex ratios of pups, and pup body weights on PND5 (Figure 5E).

## DISCUSSION

In the present study, F0 females were exposed to cannabis vapor at 8 am and 5 pm (2x/day) with doses of 25, 100, or 200 mg/ml THC of cannabis extract during pregnancy and nursing. Plasma THC concentrations in F0 females reached 25-50 ng/ml from doses of 25-200 mg/ml THC immediately after the last cannabis exposure. While maximum THC levels in maternal plasma (Cmax) cannot be measured in humans for obvious reasons, Kumar *et al*. have reported predicting such levels in maternal and/or fetal THC levels using their model system, maternal-fetal physiologically based pharmacokinetic (m-f-PBPK). The m-f PBPK model predicts that 10 mg of daily THC inhalation can reach over 30 ng/ml in maternal plasma, which is similar to non-pregnant women (Kumar et al. 2025). THC-COOH is known as a major metabolite of THC that has a longer half-life and is generally used for the detection of cannabis use in humans. Plasma levels of THC-COOH in humans have been reported to be 7-60 ng/ml in frequent cannabis smoke users (Karschner et al. 2009; Schwope et al. 2011). Our F0 females showed 10-20 ng/ml of THC-COOH in plasma. Thus, those levels were biologically relevant to human exposure levels in chronic users. Importantly, until recently, studies describing the effects of cannabis have relied on administering THC to rodents via injection or oral gavage (Asch and Smith 1986; de Salas-Quiroga et al. 2015; Natale et al. 2020). However, inhalation, such as smoking and vaporizing, is the most common route of cannabis administration among pregnant women (Young-Wolff et al. 2022; Young-Wolff et al. 2021). Therefore, vaporization is an ideal administration method to study the effect of cannabis on reproductive function.

Our study showed that cannabis exposure during the entire pregnancy did not induce preterm birth or stillbirth. We did not observe any health issues in F0 females, either. While many reports indicate the association between cannabis use and an increased risk of preterm birth (Conner et al. 2016; Duko et al. 2023; Gunn et al. 2016; Lo et al. 2024; Marchand et al. 2022), women who use cannabis during pregnancy often take other substances, such as alcohol, tobacco, and/or other illicit drugs (Chabarria et al. 2016). Features of cannabis use in pregnancy include being single or unmarried or having a lower income and socioeconomic status (Fried et al. 1980). We thus cannot directly extrapolate these results to human cannabis users because of the myriad factors that are associated with cannabis-related outcomes in humans. However, our data can provide more specific insight into the effects of cannabis alone, independent of all of these extraneous factors. We should also consider that mice can have multiple pups at once, as shown by having more than 10 pups in F0 CD1 female mice. In this study, F1 pup numbers were not affected by in utero cannabis exposure. In support of this, there is limited inconclusive evidence to suggest the association between prenatal cannabis exposure and stillbirth (Conner et al. 2016; Gunn et al. 2016; Lo et al. 2024; Marchand et al. 2022). In contrast, neonatal body weights were smaller in cannabis exposed F1 offspring. This result supports the risks of reduced fetal growth, low birth weight, and increased neonatal intensive care by prenatal cannabis exposure (Fonseca and Rebelo 2021; Hayer et al. 2023; Lo et al. 2022).

Longitudinal studies have recently reported an increased association between maternal cannabis use and adverse effects on fetal and neonatal developmental outcomes (Gunn et al. 2016). These studies suggest an association between maternal cannabis use and adverse cognitive and behavioral outcomes in offspring (Day et al. 1992; El Marroun et al. 2009; Fried et al. 1984), possibly due to disruption of proper brain development (Brummelte et al. 2017; Nashed et al. 2020; Wu et al. 2011). In the present study, perinatal cannabis exposure delayed puberty onset and disrupted estrous cyclicity. Although there is no clear evidence that maternal cannabis use leads to delayed puberty, our results suggest that the hypothalamus and/or pituitary could be affected by maternal cannabis exposure, as GnRH in the hypothalamus and LH/FSH in the pituitary are crucial to trigger the stimulation of ovary for puberty onset. THC can interfere with the ECS in the hypothalamus and pituitary via the CB1 receptor (Ryan et al. 2021). Prenatal THC exposure can interfere with folliculogenesis and reduce ovarian vasculature in rats (Martinez-Pena et al. 2021). However, cannabis-exposed ovarian and uterine function in adults was normal, showing no alterations of plasma estradiol-17β levels, ovarian and uterine histology, and pregnancy outcomes. Indeed, body weight, as well as ovarian weight, was within a normal range by the time they reached adulthood, indicating that maternal cannabis exposure did not consistently affect offspring growth after weaning. Notably, this could help to explain why no phenotypic reproductive defects were observed in cannabis-exposed F1 adult females.

The mechanisms of how cannabis affects the health outcomes for the next generation may occur via epigenetic modifications. Recent studies have demonstrated that parental THC exposure (Szutorisz et al. 2014; Watson et al. 2015) and paternal exposure to the synthetic CB1 receptor agonist WIN55,212-2 elicits phenotypic behavior impairment correlated with altered DNA methylomes in the brain of F1 rat offspring (Ibn Lahmar Andaloussi et al. 2019). A study from Murphy’s group highlights the ability of THC exposure to induce significant changes in DNA methylomes in rat sperm (Murphy et al. 2018). Many “THC target genes” are differentially methylated in both rat sperm and brain (Murphy et al. 2018; Watson et al. 2015), suggesting that reproductive risks could be associated with abnormal behavior via epigenetic modification in sperm. This group has further demonstrated that DNA methylomes in human sperm are altered in cannabis users compared with those in non-user controls (Murphy et al. 2018). Thus, germ cell epimutation can be transmitted to the next generations to induce phenotypic defects in offspring. As no rigorous screening of maternal germline transmission to characterize phenotypic defects caused by maternal cannabis exposure has been performed, we examined additional generations: F2 as multigenerational (directly exposed) and F3 as transgenerational (indirectly exposed). However, phenotypic female reproductive parameters were not affected in maternal cannabis-exposed F2 and F3 offspring, although estrous cyclicity was disrupted in the developmental stage of F2 females. These results suggest that the transgenerational effect of maternal cannabis exposure is limited, and defects induced by cannabis are less likely to cause the disruption of female reproductive function across generations. Nevertheless, altered DNA methylation and/or epigenetic modification caused by cannabis exposure in oogenesis remains to be studied.

Collectively, the present study showed that perinatal cannabis vapor exposure did not impair pregnancy in F0 females but disrupted developmental stages of female reproduction, mainly in F1 offspring. In contrast, perinatal cannabis exposure did not affect adult female reproductive function in F1, F2, and F3 offspring. Our results indicate that the transgenerational effect of cannabis vapor exposure on female reproduction is limited, and reproductive alterations caused by cannabis and their mechanisms might be exclusive to the directly exposed generation.

## Author contributions

M.S. and K.H. designed the research; M.S., Y.O., and D.A.M. performed research and analyzed data; J.A.M. and R.J.M. provided critical feedback on the manuscript; K.H. wrote the paper; all authors read, reviewed, edited, and approved the manuscript.

## REFERENCES

(SAMHSA) SAaMHSA. 2020. 2019 national survey on drug use and health (nsduh) annual national report. U.S. Department of Health & Human Services.

American College of O, Gynecologists Committee on Obstetric P. 2015. Committee opinion no. 637: Marijuana use during pregnancy and lactation. Obstet Gynecol. 126(1):234–238.

Asch RH, Smith CG. 1986. Effects of delta 9-thc, the principal psychoactive component of marijuana, during pregnancy in the rhesus monkey. J Reprod Med. 31(12):1071–1081.

Baggio S, Deline S, Studer J, Mohler-Kuo M, Daeppen JB, Gmel G. 2014. Routes of administration of cannabis used for nonmedical purposes and associations with patterns of drug use. J Adolesc Health. 54(2):235–240.

Belladelli F, Del Giudice F, Kasman A, Kold Jensen T, Jorgensen N, Salonia A, Eisenberg ML. 2021. The association between cannabis use and testicular function in men: A systematic review and meta-analysis. Andrology. 9(2):503–510.

Blackard C, Tennes K. 1984. Human placental transfer of cannabinoids. N Engl J Med. 311(12):797.

Brents LK. 2016. Marijuana, the endocannabinoid system and the female reproductive system. Yale J Biol Med. 89(2):175–191.

Britch SC, Wiley JL, Yu Z, Clowers BH, Craft RM. 2017. Cannabidiol-delta(9)- tetrahydrocannabinol interactions on acute pain and locomotor activity. Drug Alcohol Depend. 175:187–197.

Brummelte S, Mc Glanaghy E, Bonnin A, Oberlander TF. 2017. Developmental changes in serotonin signaling: Implications for early brain function, behavior and adaptation. Neuroscience. 342:212–231.

Carvalho RK, Andersen ML, Mazaro-Costa R. 2020. The effects of cannabidiol on male reproductive system: A literature review. J Appl Toxicol. 40(1):132–150.

Chabarria KC, Racusin DA, Antony KM, Kahr M, Suter MA, Mastrobattista JM, Aagaard KM. 2016. Marijuana use and its effects in pregnancy. Am J Obstet Gynecol. 215(4):506 e501–507.

Chakravarty I, Shah PG, Sheth AR, Ghosh JJ. 1979. Mode of action of delta-9- tetrahydrocannabinol on hypothalamo-pituitary function in adult female rats. J Reprod Fertil. 57(1):113–115.

Chandra S, Radwan MM, Majumdar CG, Church JC, Freeman TP, ElSohly MA. 2019. New trends in cannabis potency in USA and europe during the last decade (2008-2017). Eur Arch Psychiatry Clin Neurosci. 269(1):5–15.

Conner SN, Bedell V, Lipsey K, Macones GA, Cahill AG, Tuuli MG. 2016. Maternal marijuana use and adverse neonatal outcomes: A systematic review and meta-analysis. Obstet Gynecol. 128(4):713–723.

Day N, Cornelius M, Goldschmidt L, Richardson G, Robles N, Taylor P. 1992. The effects of prenatal tobacco and marijuana use on offspring growth from birth through 3 years of age. Neurotoxicol Teratol. 14(6):407–414.

Day NL, Goldschmidt L, Day R, Larkby C, Richardson GA. 2015. Prenatal marijuana exposure, age of marijuana initiation, and the development of psychotic symptoms in young adults. Psychol Med. 45(8):1779–1787.

Day NL, Leech SL, Goldschmidt L. 2011. The effects of prenatal marijuana exposure on delinquent behaviors are mediated by measures of neurocognitive functioning. Neurotoxicol Teratol. 33(1):129–136.

de Salas-Quiroga A, Diaz-Alonso J, Garcia-Rincon D, Remmers F, Vega D, Gomez-Canas M, Lutz B, Guzman M, Galve-Roperh I. 2015. Prenatal exposure to cannabinoids evokes long-lasting functional alterations by targeting cb1 receptors on developing cortical neurons. Proc Natl Acad Sci U S A. 112(44):13693–13698.

Dhawan K, Sharma A. 2003. Restoration of chronic-delta 9-thc-induced decline in sexuality in male rats by a novel benzoflavone moiety from passiflora incarnata linn. Br J Pharmacol. 138(1):117–120.

Di Blasio AM, Vignali M, Gentilini D. 2013. The endocannabinoid pathway and the female reproductive organs. J Mol Endocrinol. 50(1):R1–9.

Dubovis M, Muneyyirci-Delale O. 2020. Effects of marijuana on human reproduction. Reprod Toxicol. 94:22–30.

Duko B, Dachew BA, Pereira G, Alati R. 2023. The effect of prenatal cannabis exposure on offspring preterm birth: A cumulative meta-analysis. Addiction. 118(4):607–619.

El Marroun H, Tiemeier H, Steegers EA, Jaddoe VW, Hofman A, Verhulst FC, van den Brink W, Huizink AC. 2009. Intrauterine cannabis exposure affects fetal growth trajectories: The generation r study. J Am Acad Child Adolesc Psychiatry. 48(12):1173–1181.

ElSohly MA, Mehmedic Z, Foster S, Gon C, Chandra S, Church JC. 2016. Changes in cannabis potency over the last 2 decades (1995-2014): Analysis of current data in the united states. Biol Psychiatry. 79(7):613–619.

Fine JD, Moreau AL, Karcher NR, Agrawal A, Rogers CE, Barch DM, Bogdan R. 2019. Association of prenatal cannabis exposure with psychosis proneness among children in the adolescent brain cognitive development (abcd) study. JAMA Psychiatry. 76(7):762–764.

Fonseca BM, Rebelo I. 2021. Cannabis and cannabinoids in reproduction and fertility: Where we stand. Reprod Sci.

Freels TG, Baxter-Potter LN, Lugo JM, Glodosky NC, Wright HR, Baglot SL, Petrie GN, Yu Z, Clowers BH, Cuttler C et al. 2020. Vaporized cannabis extracts have reinforcing properties and support conditioned drug-seeking behavior in rats. J Neurosci. 40(9):1897–1908.

Fried PA, Watkinson B, Grant A, Knights RM. 1980. Changing patterns of soft drug use prior to and during pregnancy: A prospective study. Drug Alcohol Depend. 6(5):323–343.

Fried PA, Watkinson B, Willan A. 1984. Marijuana use during pregnancy and decreased length of gestation. Am J Obstet Gynecol. 150(1):23–27.

Goldschmidt L, Day NL, Richardson GA. 2000. Effects of prenatal marijuana exposure on child behavior problems at age 10. Neurotoxicol Teratol. 22(3):325–336.

Grant KS, Petroff R, Isoherranen N, Stella N, Burbacher TM. 2018. Cannabis use during pregnancy: Pharmacokinetics and effects on child development. Pharmacol Ther. 182:133–151.

Gunn JK, Rosales CB, Center KE, Nunez A, Gibson SJ, Christ C, Ehiri JE. 2016. Prenatal exposure to cannabis and maternal and child health outcomes: A systematic review and meta-analysis. BMJ Open. 6(4):e009986.

Hayer S, Mandelbaum AD, Watch L, Ryan KS, Hedges MA, Manuzak JA, Easley CAt, Schust DJ, Lo JO. 2023. Cannabis and pregnancy: A review. Obstet Gynecol Surv. 78(7):411–428.

Ibn Lahmar Andaloussi Z, Taghzouti K, Abboussi O. 2019. Behavioural and epigenetic effects of paternal exposure to cannabinoids during adolescence on offspring vulnerability to stress. Int J Dev Neurosci. 72:48–54.

Imtiaz S, Wells S, Rehm J, Hamilton HA, Nigatu YT, Wickens CM, Jankowicz D, Elton-Marshall T. 2021. Cannabis use during the covid-19 pandemic in canada: A repeated cross-sectional study. J Addict Med. 15(6):484–490.

Karschner EL, Schwilke EW, Lowe RH, Darwin WD, Herning RI, Cadet JL, Huestis MA. 2009. Implications of plasma delta9-tetrahydrocannabinol, 11-hydroxy-thc, and 11-nor-9- carboxy-thc concentrations in chronic cannabis smokers. J Anal Toxicol. 33(8):469–477.

Ko JY, Farr SL, Tong VT, Creanga AA, Callaghan WM. 2015. Prevalence and patterns of marijuana use among pregnant and nonpregnant women of reproductive age. Am J Obstet Gynecol. 213(2):201 e201–201 e210.

Kumar AR, Benson LS, Wymore EM, Phipers JE, Dempsey JC, Cort LA, Unadkat JD. 2025. Quantification and prediction of human fetal (-)-delta(9)-tetrahydrocannabinol/(+/−)-11- oh-delta(9)-tetrahydrocannabinol exposure during pregnancy to inform fetal cannabis toxicity. Nat Commun. 16(1):824.

Lee DC, Crosier BS, Borodovsky JT, Sargent JD, Budney AJ. 2016. Online survey characterizing vaporizer use among cannabis users. Drug Alcohol Depend. 159:227–233.

Lo JO, Hedges JC, Girardi G. 2022. Impact of cannabinoids on pregnancy, reproductive health, and offspring outcomes. Am J Obstet Gynecol. 227(4):571–581.

Lo JO, Shaw B, Robalino S, Ayers CK, Durbin S, Rushkin MC, Olyaei A, Kansagara D, Harrod CS. 2024. Cannabis use in pregnancy and neonatal outcomes: A systematic review and meta-analysis. Cannabis Cannabinoid Res. 9(2):470–485.

Marchand G, Masoud AT, Govindan M, Ware K, King A, Ruther S, Brazil G, Ulibarri H, Parise J, Arroyo A et al. 2022. Birth outcomes of neonates exposed to marijuana in utero: A systematic review and meta-analysis. JAMA Netw Open. 5(1):e2145653.

Martinez-Pena AA, Lee K, Petrik JJ, Hardy DB, Holloway AC. 2021. Gestational exposure to delta(9)-thc impacts ovarian follicular dynamics and angiogenesis in adulthood in wistar rats. J Dev Orig Health Dis. 12(6):865–869.

Mendelson JH, Mello NK, Ellingboe J. 1985. Acute effects of marihuana smoking on prolactin levels in human females. J Pharmacol Exp Ther. 232(1):220–222.

Murphy SK, Itchon-Ramos N, Visco Z, Huang Z, Grenier C, Schrott R, Acharya K, Boudreau MH, Price TM, Raburn DJ et al. 2018. Cannabinoid exposure and altered DNA methylation in rat and human sperm. Epigenetics. 13(12):1208–1221.

Nashed MG, Hardy DB, Laviolette SR. 2020. Prenatal cannabinoid exposure: Emerging evidence of physiological and neuropsychiatric abnormalities. Front Psychiatry. 11:624275.

Natale BV, Gustin KN, Lee K, Holloway AC, Laviolette SR, Natale DRC, Hardy DB. 2020. Delta9-tetrahydrocannabinol exposure during rat pregnancy leads to symmetrical fetal growth restriction and labyrinth-specific vascular defects in the placenta. Sci Rep. 10(1):544.

Pertwee RG, Howlett AC, Abood ME, Alexander SP, Di Marzo V, Elphick MR, Greasley PJ, Hansen HS, Kunos G, Mackie K et al. 2010. International union of basic and clinical pharmacology. Lxxix. Cannabinoid receptors and their ligands: Beyond cb(1) and cb(2). Pharmacol Rev. 62(4):588–631.

Porath AJ, Fried PA. 2005. Effects of prenatal cigarette and marijuana exposure on drug use among offspring. Neurotoxicol Teratol. 27(2):267–277.

Reece AS, Hulse GK. 2019. Cannabis teratology explains current patterns of coloradan congenital defects: The contribution of increased cannabinoid exposure to rising teratological trends. Clin Pediatr (Phila). 58(10):1085–1123.

Ryan KS, Mahalingaiah S, Campbell LR, Roberts VHJ, Terrobias JJD, Naito CS, Boniface ER, Borgelt LM, Hedges JC, Hanna CB et al. 2021. The effects of delta-9- tetrahydrocannabinol exposure on female menstrual cyclicity and reproductive health in rhesus macaques. F S Sci. 2(3):287–294.

Schwope DM, Karschner EL, Gorelick DA, Huestis MA. 2011. Identification of recent cannabis use: Whole-blood and plasma free and glucuronidated cannabinoid pharmacokinetics following controlled smoked cannabis administration. Clin Chem. 57(10):1406–1414.

Shi M, Sekulovski N, MacLean JA, 2nd, Hayashi K. 2017. Effects of bisphenol a analogues on reproductive functions in mice. Reprod Toxicol. 73:280–291.

Shi M, Sekulovski N, MacLean JA, Whorton A, Hayashi K. 2019a. Prenatal exposure to bisphenol a analogues on female reproductive functions in mice. Toxicol Sci. 168(2):561–571.

Shi M, Whorton AE, Sekulovski N, MacLean JA, Hayashi K. 2019b. Prenatal exposure to bisphenol a, e, and s induces transgenerational effects on female reproductive functions in mice. Toxicol Sci. 170(2):320–329.

Sonon KE, Richardson GA, Cornelius JR, Kim KH, Day NL. 2015. Prenatal marijuana exposure predicts marijuana use in young adulthood. Neurotoxicol Teratol. 47:10–15.

Szutorisz H, DiNieri JA, Sweet E, Egervari G, Michaelides M, Carter JM, Ren Y, Miller ML, Blitzer RD, Hurd YL. 2014. Parental thc exposure leads to compulsive heroin-seeking and altered striatal synaptic plasticity in the subsequent generation. Neuropsychopharmacology. 39(6):1315–1323.

van Gelder MM, Donders AR, Devine O, Roeleveld N, Reefhuis J, National Birth Defects Prevention S. 2014. Using bayesian models to assess the effects of under-reporting of cannabis use on the association with birth defects, national birth defects prevention study, 1997-2005. Paediatr Perinat Epidemiol. 28(5):424–433.

van Gelder MM, Reefhuis J, Caton AR, Werler MM, Druschel CM, Roeleveld N, National Birth Defects Prevention S. 2010. Characteristics of pregnant illicit drug users and associations between cannabis use and perinatal outcome in a population-based study. Drug Alcohol Depend. 109(1-3):243–247.

Volkow ND, Han B, Compton WM, McCance-Katz EF. 2019. Self-reported medical and nonmedical cannabis use among pregnant women in the united states. JAMA. 322(2):167–169.

Walker OS, Holloway AC, Raha S. 2019. The role of the endocannabinoid system in female reproductive tissues. J Ovarian Res. 12(1):3.

Watson CT, Szutorisz H, Garg P, Martin Q, Landry JA, Sharp AJ, Hurd YL. 2015. Genome-wide DNA methylation profiling reveals epigenetic changes in the rat nucleus accumbens associated with cross-generational effects of adolescent thc exposure. Neuropsychopharmacology. 40(13):2993–3005.

Weinsheimer RL, Yanchar NL, Canadian Pediatric Surgical N. 2008. Impact of maternal substance abuse and smoking on children with gastroschisis. J Pediatr Surg. 43(5):879–883.

Williams LJ, Correa A, Rasmussen S. 2004. Maternal lifestyle factors and risk for ventricular septal defects. Birth Defects Res A Clin Mol Teratol. 70(2):59–64.

Wu CS, Jew CP, Lu HC. 2011. Lasting impacts of prenatal cannabis exposure and the role of endogenous cannabinoids in the developing brain. Future Neurol. 6(4):459–480.

Young-Wolff KC, Adams SR, Brown QL, Weisner C, Ansley D, Goler N, Skelton KR, Satre DD, Foti TR, Conway A. 2022. Modes of cannabis administration in the year prior to conception among patients in northern california. Addict Behav Rep. 15:100416.

Young-Wolff KC, Ray GT, Alexeeff SE, Adams SR, Does MB, Ansley D, Avalos LA. 2021. Rates of prenatal cannabis use among pregnant women before and during the covid-19 pandemic. JAMA. 326(17):1745–1747.

Zammit S, Thomas K, Thompson A, Horwood J, Menezes P, Gunnell D, Hollis C, Wolke D, Lewis G, Harrison G. 2009. Maternal tobacco, cannabis and alcohol use during pregnancy and risk of adolescent psychotic symptoms in offspring. Br J Psychiatry. 195(4):294–300.

